# An unsupervised framework for comparing SARS-CoV-2 protein sequences using LLMs

**DOI:** 10.1101/2024.12.16.628708

**Authors:** Sayantani B. Littlefield, Roy H. Campbell

## Abstract

The severe acute respiratory syndrome coronavirus 2 (SARS-CoV-2) pandemic led to 700 million infections and 7 million deaths worldwide. While studying these viruses, scientists developed a large amount of sequencing data that was made available to researchers. Large language models (LLMs) are pre-trained on large databases of proteins and prior work has shown its use in studying the structure and function of proteins. This paper proposes an unsupervised framework for characterizing SARS-CoV-2 sequences using large language models. First, we perform a comparison of several language models previously proposed by other authors. This step is used to determine how clustering and classification approaches perform on SARS-CoV-2 sequence embeddings. In this paper, we focus on surface glycoprotein sequences, also known as spike proteins in SARS-CoV-2 because scientists have previously studied their involvement in being recognized by the human immune system. Our contrastive learning framework is trained in an unsupervised manner, leveraging the Levenshtein distance from pairwise alignment of sequences when the contrastive loss is computed by the Siamese Neural Network. The final part of this paper focuses on a comparison with a previous approach on a test dataset containing data from the latter part of the pandemic. In the prediction of emerging variants, the proposed LLM-based approach shows an improvement of 0.2 in terms of the adjusted rand index clustering compared to a previously proposed approach. This shows the potential of applying large language models to this field.

## 1 Introduction

The evolution of pandemics like coronavirus disease 2019 (COVID-19) have caused widespread infection and deaths worldwide [1]. Pandemics typically involve health officials introducing a set of mitigation measures while researchers develop ways to study vaccines and design drugs for the benefit of society [2]. While studying this virus, prior work has shown there has been a deluge of sequencing data from pandemics. This information is stored publicly by the National Institute of Heath-the National Center of Biotechnology Information (https://www.ncbi.nlm.nih.gov). Computational techniques are valuable in the detection of SARS-CoV-2 variants [3, 4, 5, 6].

In this paper, we applied protein language models (pLMs) to studying SARS-CoV-2 surface glycoprotein sequences because they are recognized by the human immune system [7, 8]. pLMs are a set of models applicable to protein sequences, which generate condensed representations in the form of embedding vectors. Similarly, we applied genome language models (gLMs) to studying SARS-CoV-2 gene sequences. gLMs are trained on genomes and are capable of identifying long-range interactions and regulatory elements [9]. A comparison study on chronologically ordered SARS-CoV-2 sequences will help understand the performance of these embedding vectors relative to the variant labels [10].

Following our comparison study, we extend our previously proposed framework for supervised contrastive learning to unsupervised conditions. Contrastive learning is used in machine learning to identify differences among different samples. Such a comparison will eventually help with identification of clusters of variants [11]. We hypothesize that similarities in protein sequences that are reflected in their corresponding n-dimensional LLM sequence embeddings can be used to train a contrastive framework and observe similarities and differences in protein sequences.

The rest of this paper is structured as follows: Section 2 provides an overview of related work, Section 3 discusses the datasets used, Section 4 covers the methods used in our comparison study and unsupervised contrastive learning framework, Section 5 discusses the results obtained, and Section 6 concludes the paper.

## 2 Related Work

This section provides a review of existing literature in methods used for analysis of variant data. Computational software like Nextalign and Nextclade [12, 13, 14] applied to labeling SARS-CoV-2 variants rely on sequence alignment, which is computationally expensive. Approaches involving k-mers where k residues are used to form ‘words’ from a protein sequence to allow for an alignment-free approach. While prior work has also shown that k-mer based approaches perform better than alignment based approaches [15], they are subject to erroneous associations and also lead to high computational complexity in certain cases.

Several other machine learning approaches have been proposed on SARS-CoV-2 data, starting with classification of coronaviruses at various levels of taxa [16, 17]. Brierley and Fowler [18] classified intermediate COVID-19 hosts with random forests where the models were trained on SARS-CoV-2 spike proteins and gene sequences. The authors claimed deriving “evolutionary signatures” from genomic data may be possible, and this could indicate species origin [10].

Basu and Campbell [19] previously proposed a long short-term memory (LSTM) deep learning model that enabled the classification of different SARS-CoV-2 variants. However, the model was trained in 2021 on an early COVID-19 snapshot of twenty variants at the time. The structure of the proposed network required re-training each time newer SARS-CoV-2 variants added to the database.

ProtVec [20] and ProteinBERT [21] were language models in the field before the introduction of large language models. ProtVec is inspired by word2vec [22]. The model was pre-trained on 546,790 SwissProt sequences. It uses a skip-gram neural network to train positive and negative 3-mers depending on the context. ProteinBERT [21] is an influential paper in this field modeled after BERT [23]. It consists of six blocks of transformers as well as fully connected and convolutional layers. The global attention layers exhibit linear complexity with respect to sequence length, which makes the model adaptable to sequences of different lengths. This model was pre-trained on 106M UniRef90 proteins. Large language models (LLMs) gained popularity in natural language applications with the introduction of transformers [24, 23] and generative pre-trained transformers (GPT) [25]. A lot of these models have gained popularity in exploring viral fitness and predicting viral escape [26, 27].

We start with looking at a set of protein language models (pLMs). We compare the performance of a set of previously published protein language models ProtVec [20], ProteinBERT [21], ESM-1v and ESM-2 [28] in order to determine the classification and clustering performance on SARS-CoV-2 spike protein sequences. Evolutionary scale modeling (ESM) consist of a set of protein transformer models. These models have a masked language model (MLM) objective which is used to determine protein structure and contact prediction using representation learning and self-attention [29]. ESM-1v and ESM-2 are the associated research models proposed by Lin et al. [28]. In addition to our work on pLMs, we also surveyed a set of genome language models (gLMs) including genome-scale language model (GenSLM) [30] uses attention to determine evolutionary dynamics as well as HyenaDNA that uses long convolutions [31]. We only discuss a subset of research models in this paper when there are several other models that exist in this field [32, 33, 34].

Contrastive learning is a strategy used in machine learning where the model learns to distinguish among samples using comparisons. Siamese Neural Networks (SNNs) are a type of contrastive learning model originally used in computer vision [35]. SNNs operate by sharing weights and learn to minimize the distance between similar classes while separating the distance between different classes. In the field of viral genomics, SNN literature is limited. Madan et al. [36] used SNNs on spike protein embeddings to predict protein-protein interaction (PPI) networks for SARS-CoV-2. The authors applied ProteinBERT to generate the embeddings and showed how the properties of proteins can be learned from sequence data. This work is an extension of Tsukiyama et al. [37] where an (LSTM) model was applied to word2vec embeddings to study PPI interactions. From our previous work on using SNNs in supervised settings [38], we observed that SNNs have potential in differentiating various classes and choose to extend it to unsupervised settings in this paper. Our inspiration for comparison in an unsupervised manner also stems from recent similar work [39] on spike proteins that studied the detection of emerging variants in such settings.

In this paper, we first discuss a cursory comparison of various language models in a supervised manner, following which we discuss the unsupervised contrastive learning framework.

## 3 Datasets

SARS-CoV-2 surface glycoprotein and gene sequences were downloaded from the National Center for Biotechnology Information (NCBI), an NIH sequenced data repository. The sequences were filtered based on completeness (complete), type of protein (surface glycoproteins), host (*Homo sapiens*) and country (United States).

### 3.1 Pre-processing

The NCBI interface allowed to easily filter and download spike protein sequences. Similar to the protein sequences, focusing on the spike region of the gene sequences enabled a fair comparison when applying language models. This also helped from the point of data storage since SARS-CoV-2 whole genome sequences each were 29,000 base pairs on average, making it time-consuming to focus on portions from the entire genome that would not appropriately correspond to the spike protein sequences. The following process was used to extract the spike genome portions from the full-length genomes:

1. The positions of the reference SARS-CoV-2 (Wuhan wildtype) are known to be 21563 to 25384 where the indices are 1-indexed and both are inclusive. This means the spike genome in the reference wildtype sequence has a total of 25384 *−* 21563 + 1 = 3822 base pairs.
2. Now that the reference spike genome indices are known, the full-length genomes were then aligned pair-wise with minimap2 with the asm5 flag that enables aligning assemblies that are highly similar with 0-5% divergence.
3. Finally, the gaps resulting from pairwise alignment in the previous step are removed to obtain the spike genome sequences.

### 3.2 Supervised comparison study

The pango nomenclature lineage information was extracted from the same NCBI dataset, while the lineage classification (Omicron, Alpha, Beta, Delta, Gamma) was extracted from the https://cov-lineages.org [40, 41]. The cov-lineages dataset had information about the individual lineages and which variants belonged to each lineage. According to the website, B.1.1.529, BA.1, BA.2, BA.5, XBB, XBB.1, XBB.1.5, and XBB.1.6 belong to the Omicron lineage, B.1.1.7 belongs to the Alpha lineage, B.1.351 belongs to the Beta lineage, B.1.617.2 belongs to the Delta lineage, and P.1 belongs to the Gamma lineage. The dataset (excluding the held-out set for emergent variant analysis) was divided into 80% train and 20% validation data with random state=0 and stratification to enable a fair supervised comparison across all models with non-zero class frequencies for all five lineages.

### 3.3 Unsupervised contrastive learning approach

Following pre-processing from the previous subsection, these were chronologically ordered records with PANGO labels extracted from NCBI. The entire set of chronologically ordered records had sample collection dates from January 2020 to December 2025. The unsupervised contrastive learning approach required the chronological ordering to allow the model to predict previously unseen variants over time. For reproducibility, it was split chronologically into 80% (1,145,700 sequences) and 20% (286,425 sequences) as the train + validation and test sets (held-out set for emergent variant analysis) respectively.

## 4 Methods

In this section, we introduce the methods used in the supervised comparison study and the unsupervised contrastive learning approach.

### 4.1 Comparison Study

Surface glycoprotein sequences were obtained from NCBI Virus (NIH), a public repository of sequencing data. Variant labels were extracted from the same NCBI Virus dataset and linked to the lineage classification from https://cov-lineages.org [40, 41]. We extracted five main lineages: Omicron (B.1.1.529, BA.1, BA.2, BA.5, XBB, XBB.1, XBB.1.5, and XBB.1.6), Alpha (B.1.1.7), Beta (B.1.351), Delta (B.1.617.2), and Gamma (P.1). The labels for this supervised study are assigned based on the lineages.

### 4.2 Contrastive Learning Framework

Contrastive learning models learn by comparing existing classes. Siamese Neural Networks (SNNs) [42, 35] were originally used in computer vision to identify new classes at the inference step based on finding similarity among previously existing classes. We consider an SNN for the contrastive learning framework as previous literature has indicated that many bioinformatics tools currently struggle with the introduction of novel viral sequences [43]. Developing such a tool is useful in the case of viral pandemic data where new variants are on the rise, as it has the ability to handle newer sequences and variants after the pipeline is already trained on data from previously circulating variants. SNNs operate by sharing weights that minimize the distance between similar classes, while separating the distance between different classes. This changes the embeddings in the n-dimensional space while training the network, which we observe clarifies the clusters as indicated by cluster metrics.

Following our comparison study, we extend our previously proposed Siamese Neural Network (SNN) framework [38] for supervised contrastive learning to unsupervised conditions. The model architecture is shown in Figure 3, which shows the mirrored networks with shared weights. The objective function here is to minimize the contrastive loss. We retain the same architecture as empirically determined in our previous approach (Basu et al. [38]) for the rest of the experiments. Each network contains three convolutional layers interleaved with maxpool layers. For training the network, a batch size of 256 and a learning rate of 0.0001 were used with Adam as the optimizer [44].

All code was implemented using Python on a single node on DeltaAI and HAL [45] and later DeltaAI [46, 47].

## 5 Results and Discussion

### 5.1 Comparative analysis

In this section, we compare various language models like GenSLM [30] and ESM-2 [28] along with models like ProtVec [20] and ProteinBERT [21] on classifying and clustering pandemic viruses like that of SARS-CoV-2 into different lineages. The following metrics are used for this comparison:

- Silhouette Coefficient (SC): This metric measures similarity of points within a cluster and compares it to points belonging to different clusters. It ranges from −1 to +1. More positive values of SC indicate well-clustered data while lower values indicate poorly clustered data. It is represented as *SC* = (*b − a*)*/max*(*a, b*), where *a* is the mean intra cluster distance and *b* is the mean nearest cluster distance.
- Accuracy: This metric is computed using the number of true positives and true negatives out of all samples.
- F-1 score: This is computed as the harmonic mean of precision and recall. This ranges from 0 to 1 where higher values are preferred. The two metrics macro F-1 and weighted F-1 are represented here as Macro F-1 and weighted F-1 are represented using the following equations:

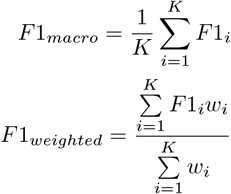

where *K* is the total number of classes and *w* is the *support* for each class. In this case, *support* for each class is defined as the number of labels belonging to the ground-truth labels for that particular class. Macro-F1 treats all classes with equal importance and generally has lower values for cases where the classifier did poorly on a specific class. Weighted F-1 has higher values even when the classifier does poorly on any specific class, but takes into account imbalanced data.

For the supervised classification task, a multi-layered Dense network, Adam optimizer [44], and a batch size of 512 is used to evaluate the performance of the models. There were five labels for SARS-CoV-2 based on the corresponding lineages. Training used EarlyStopping and the validation loss was monitored with a patience parameter of 10. The comparison study shows the accuracy and F-1 on the validation data. The macro F-1, weighted F-1, and accuracy measures are used to compare the various models. The metrics for all models and their flavors (ProtVec [20], ProteinBERT [21], GenSLM [30], ESM-1v, and ESM-2 [28], GenSLM, and HyenaDNA) are reported in Table 1. The ESM-2 model flavors (8M, 35M, 150M, 650M, and 3B) and GenSLM model flavors (25M, 250M, and 2.5B) all report the number of parameters while HyenaDNA is designed based on maximum sequence length (32k) in their published pre-trained models. Due to limits on computational resources, we did not experiment with the ESM-2 15B and GenSLM 25B models.

**Table 1:**
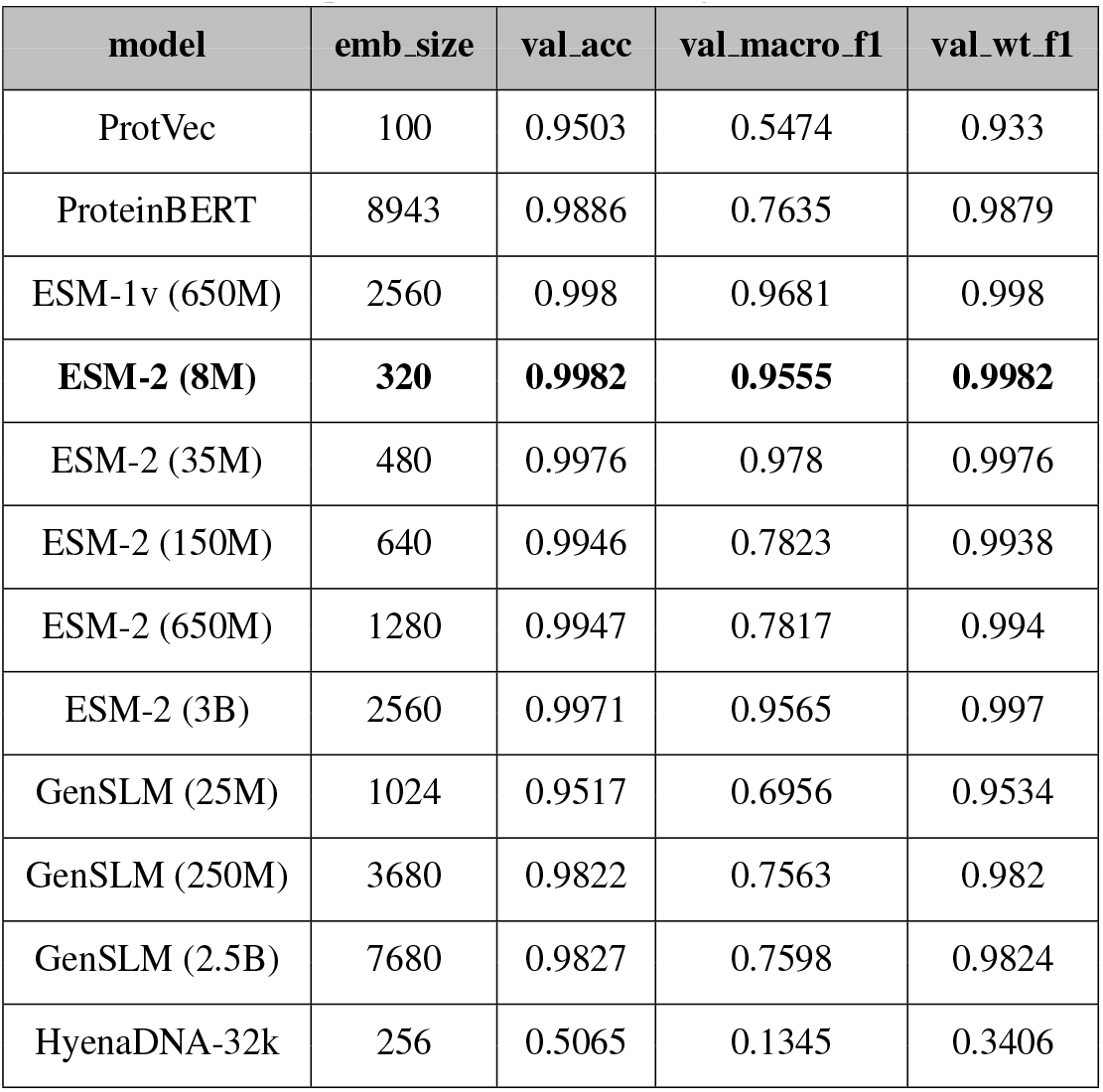
Comparison of LM embeddings on classification.

Computations were performed on the original embeddings, and t-SNE [48, 49] was only used to reduce the different embeddings to two dimensions for visualization purposes.

The results in Table 1 and Table 2 show that:

**Table 2:** Comparison of LM embeddings on clustering.

| model | emb_size | SC |
| --- | --- | --- |
| ProtVec | 100 | 0.894 |
| ProteinBERT | 8943 | 0.7829 |
| ESM-1v (650M) | 2560 | 0.8114 |
| ESM-2 (8M) | 320 | **0.8229** |
| ESM-2 (35M) | 480 | 0.7786 |
| ESM-2 (150M) | 640 | 0.8528 |
| ESM-2 (650M) | 1280 | 0.7683 |
| ESM-2 (3B) | 2560 | 0.7587 |
| GenSLM (25M) | 1024 | 0.844 |
| GenSLM (250M) | 3680 | 0.9338 |
| GenSLM (2.5B) | 7680 | 0.89 |
| HyenaDNA-32k | 256 | 0.6593 |

1. ESM-2 has the best performance on spike protein sequences when classifying among five different classes of SARS-CoV-2, with the best performing model being the 8M flavor. This was specifically chosen by looking at the validation accuracy and validation weighted F1 metrics which are the highest for this particular model.
2. To eventually study the clustering abilities *in the absence of labels*, the embeddings alone with the ground truth labels (five lineages in this case). Based on the SC standard metrics, the highest value is 0.8229 (ESM-2 8M) while the lowest value is 0.6593. Lower values of SC indicate such clusters cannot be used as-is for identifying emerging variants, especially when variants are clustered without knowledge of PANGO labels. This in turn leads us to form an approach to further separate the classes using the proposed contrastive learning framework (Siamese Neural Network). This approach makes the clusters more distinct to study emerging variants.

Figure 1 and Figure 2 show how the performance metrics vary across the different model sizes. Interestingly, for the SARS-CoV-2 dataset considered in this paper, a drop was observed in some metrics on the ESM-2 model across different sizes while the metrics remained almost consistent for the GenSLM model sizes. The performance of ESM-2 8M is almost identical to that of ESM-3B, indicating that larger models do not always work best for certain datasets. Additionally, lower dimensions (320) for the embedding size are a bonus since it meant the analysis ran faster compared to a model like ProteinBERT which has good performance but high-dimensional embeddings (8943). Since the remainder of this paper discusses this problem as an unsupervised problem, we chose to show the unsupervised clustering across the 5 lineages using a k-means clustering model with *k* = 5.

**Figure 1:**
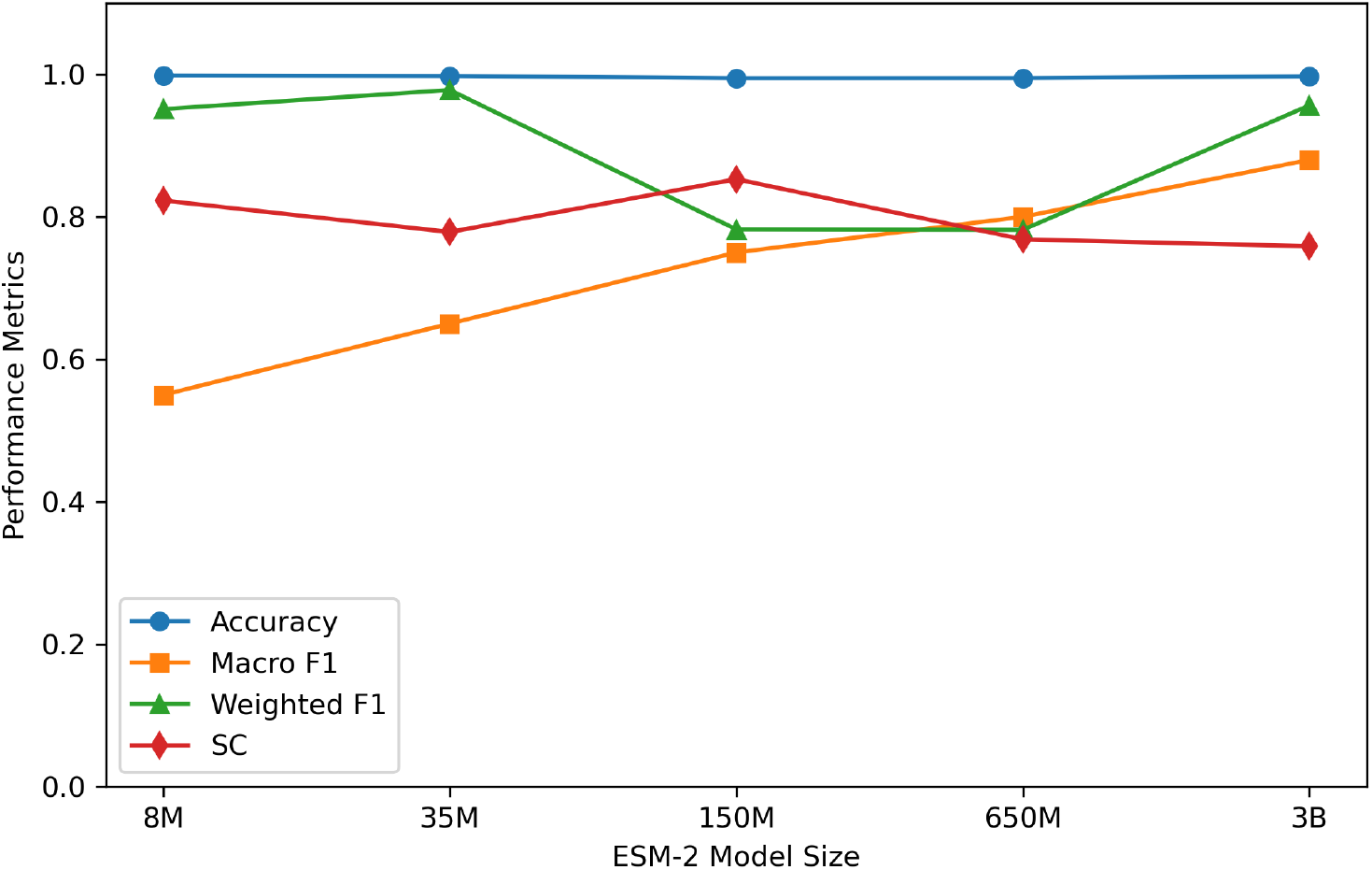
Variation of performance metrics across ESM-2 models

**Figure 2:**
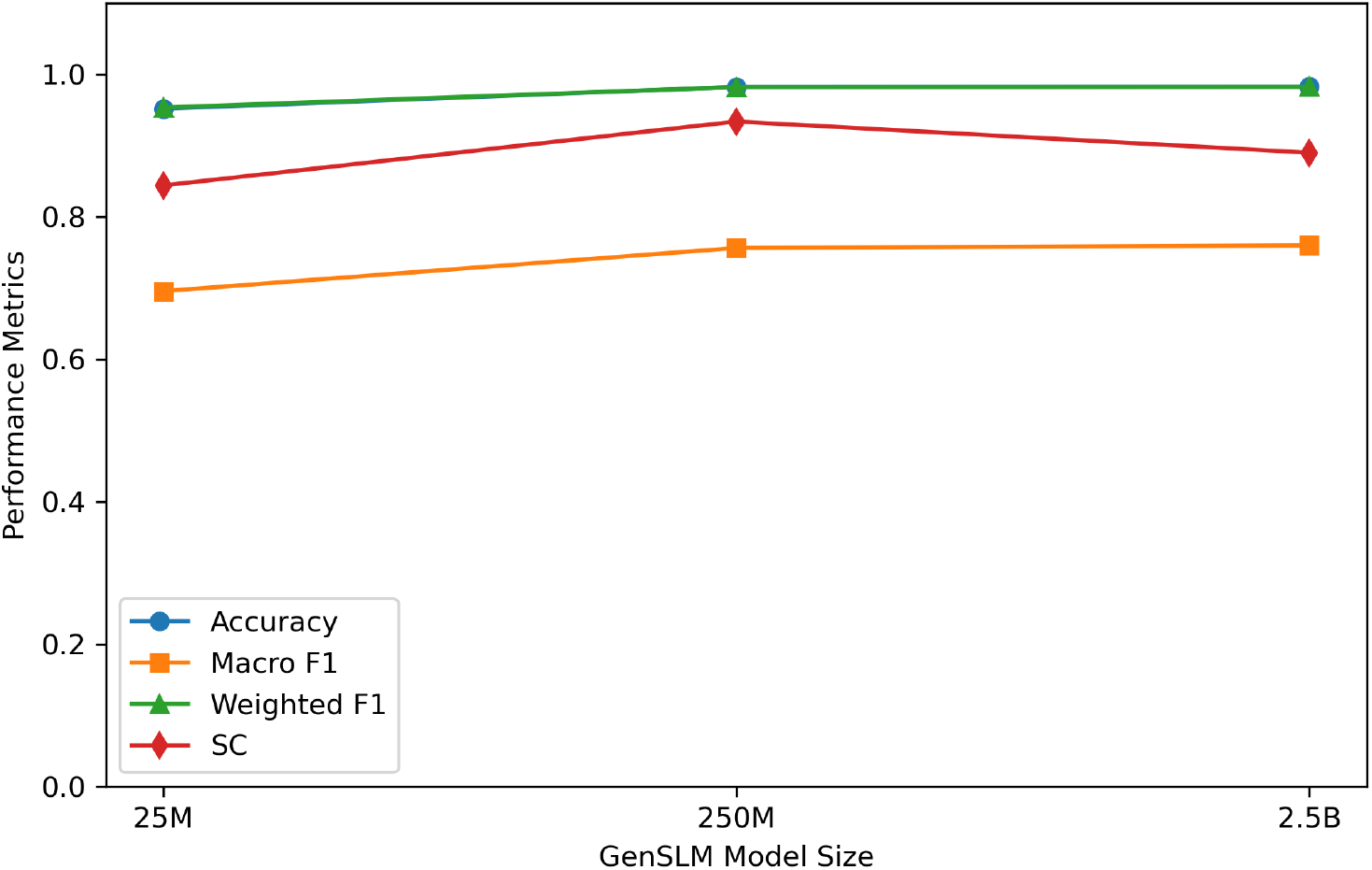
Variation of performance metrics across GenSLM models

**Figure 3:**
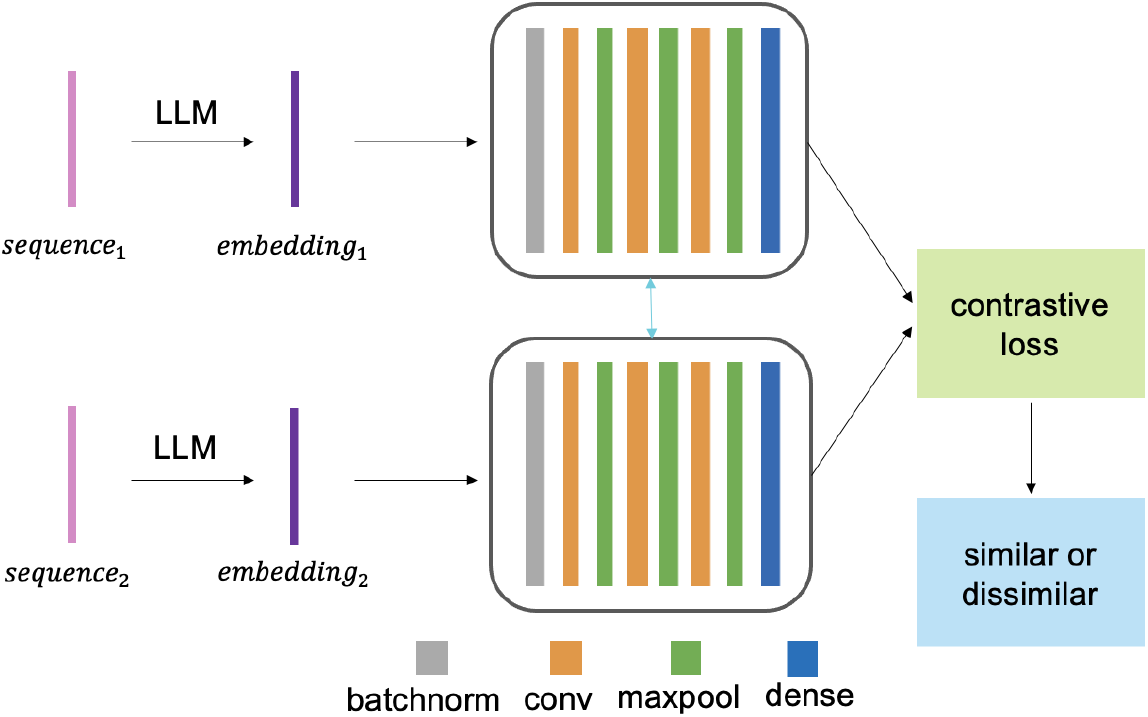
Architecture of contrastive learning framework with pairwise contrastive loss (based on Basu et al. [38])

**Figure 4:**
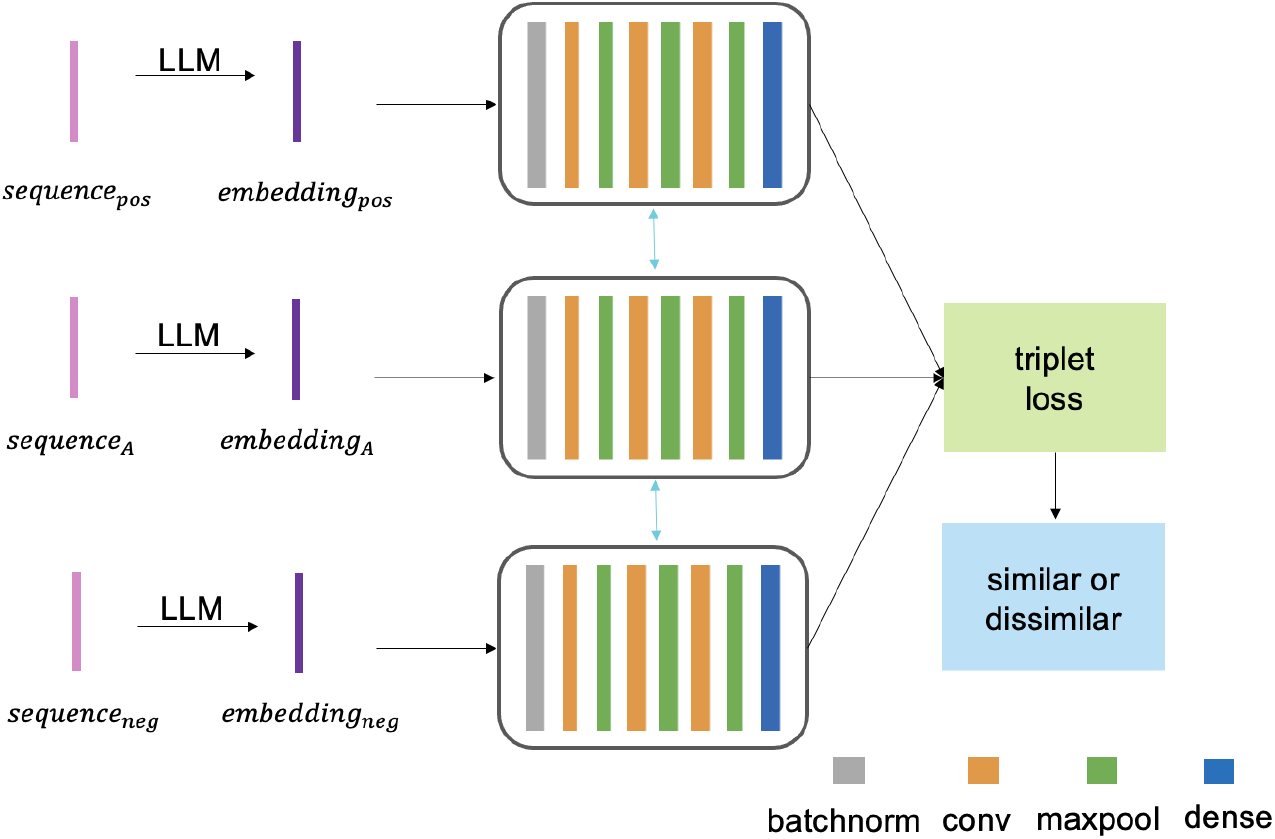
Architecture of contrastive learning framework with triplet loss (based on Basu et al. [38])

### 5.2 Unsupervised contrastive learning framework

This section details the method used in sampling the embedding pairs and triplets. All experiments were run with the ESM-2 8M embeddings from the previous section. The supervised case here is only shown as a baseline to see how much the performance drops in the unsupervised framework with the absence of labels:

1. **Supervised case:** In the supervised framework, the PANGO labels are known along with the sequences and their embeddings at the time of training. The process of mining pairs and triplets of embeddings is as follows: *Pairs:* In this case, the sequence embeddings and the corresponding PANGO labels are passed to the model. The training sample is marked similar if the two variants are the same, else they are marked dissimilar. Here, a strict comparison at the variant level is considered—for example, embedding pairs from two variants BA.1 and BA.2 are labeled ‘dissimilar’ while embedding pairs from two variants B.1.1.7 and B.1.1.7 are labeled ‘similar’. *Triplets:* In this case, three sequence embeddings and the corresponding PANGO labels are passed to the model at a time. The first embedding of the three is considered the ‘anchor’ while the remaining two are mined based on the constraint that one PANGO variant is similar to the anchor while the second PANGO variant is dissimilar to the anchor. For example, if the embedding corresponding to a sequence labeled ‘B.1.1.7’ is mined as the anchor, a corresponding positive or similar example would be an embedding corresponding to another sequence labeled ‘B.1.1.7’ while a negative or dissimilar example would be an embedding corresponding to a sequence labeled ‘BA.2’.
2. **Unsupervised case:** The unsupervised framework is more complex since the PANGO labels are not provided at the time of training the model. Only sequences and their embeddings are available at the time of training. The novelty here is using the property of Levenshtein distance to compute the similarity between the sequences using that to self-label the dataset when the LLM embeddings are provided while training. The Levenshtein distance is defined as the number of steps it takes to transform one sequence into the other. The similarity metric is then computed by calculating the percentage of positions where the residues are identical across all positions, constituting a pairwise alignment of the sequences. This metric is used to determine whether two sequences are similar or not based on two thresholds: (i) positive threshold (*pt*): if the similarity metric is higher than *pt*, the sequences are similar, and (ii) negative threshold (*nt*): if the similarity metric is lower than *nt*, the sequences are dissimilar. The process of mining pairs and triplets of embeddings is as follows: *Pairs:* In this case, the sequence embeddings and the labels computed based on the thresholds are passed to the model for training. If the similarity between the sequences is greater than *pt*, then the pair is labeled similar. If the similarity between the sequences is less than *nt*, then the pair is labeled dissimilar. *Triplets:* In this case, three sequence embeddings and the labels computed based on the thresholds are passed to the model for training. If the similarity between the positive example and the anchor is greater than *pt*, then the two embeddings are passed to the model along with the similar label. If this constraint is not satisfied, the example is skipped. If the similarity between the negative example and the anchor is lesser than *nt*, then the two embeddings are passed to the model along with the dissimilar label. If this constraint is not satisfied, the example is skipped.

Following training of the SNN, in the test phase, the SNN performance metrics are generated for the test embedding pairs. The embeddings learned by the SNN are obtained by passing the samples through one-half of the network to get the new condensed vector representations by only passing the inputs through the encoder layer. We then compared with the original ESM 8M embeddings extracted from the final layer to see how the SC metric changed.

#### Results

The chronologically ordered dataset is first split into 80% and 20% for the train and test data respectively. Note that we still have a 20% held-out test dataset from the entire dataset that we reserve to exclusively test on identifying emergent variants in the next section.

#### 5.2.1 Siamese Neural Network

Table 3 shows the complete parameter study conducted for this study varying positive threshold *pt*, negative threshold *nt* and the margin *m*. The SC is calculated using the ground truth lineage labels in both supervised and unsupervised models. This is for performance assessment to show how well the model can really perform when relying exclusively on sequence data alone. The SC for the validation set is reported at the end of each run. We used the following mathematical equations for the pairwise contrastive and triplet losses respectively, which are similar to their Tensorflow implementations, where *m* is the margin used to push dissimilar samples further apart (pairwise) or ensure the positive and negative samples are pushed further apart (triplet), *ŷ* represents the predicted value, *y* represents the ground truth (this is generated by the model in the self-supervised case where we do not provide ground-truth labels), and *N* is the number of samples.

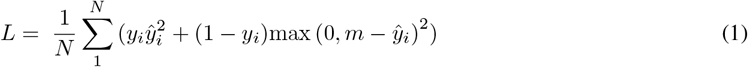

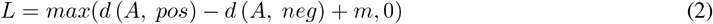

**Table 3:** Parameter study on training the proposed Siamese Neural Network (pt = positive threshold, nt = negative threshold, m = margin)

| model | loss | pt | nt | m | SC |
| --- | --- | --- | --- | --- | --- |
| unsupervised | pairwise contrastive | 0.5 | 0.5 | 0.2 | 0.7955 |
|  |  |  |  | 0.3 | 0.8085 |
|  |  |  |  | 0.4 | 0.7963 |
|  |  | 0.6 | 0.4 | 0.2 | **0.8104** |
|  |  |  |  | 0.3 | 0.789 |
|  |  |  |  | 0.4 | 0.7913 |
|  |  | 0.7 | 0.3 | 0.2 | 0.7983 |
|  |  |  |  | 0.3 | 0.8043 |
|  |  |  |  | 0.4 | 0.7981 |
|  | triplet | 0.5 | 0.5 | 0.2 | 0.7621 |
|  |  |  |  | 0.3 | 0.7717 |
|  |  |  |  | 0.4 | 0.7513 |
|  |  | 0.6 | 0.4 | 0.2 | 0.8023 |
|  |  |  |  | 0.3 | 0.768 |
|  |  |  |  | 0.4 | 0.7724 |
|  |  | 0.7 | 0.3 | 0.2 | 0.766 |
|  |  |  |  | 0.3 | 0.7562 |
|  |  |  |  | 0.4 | 0.7084 |
| supervised | pairwise contrastive | 0.5 | 0.5 | 0.2 | 0.8176 |
|  |  |  |  | 0.3 | 0.761 |
|  |  |  |  | 0.4 | 0.7776 |
|  |  | 0.6 | 0.4 | 0.2 | 0.7274 |
|  |  |  |  | 0.3 | **0.8946** |
|  |  |  |  | 0.4 | 0.7925 |
|  |  | 0.7 | 0.3 | 0.2 | 0.7951 |
|  |  |  |  | 0.3 | 0.7786 |
|  |  |  |  | 0.4 | 0.7329 |
|  | triplet | 0.5 | 0.5 | 0.2 | 0.8558 |
|  |  |  |  | 0.3 | 0.8408 |
|  |  |  |  | 0.4 | 0.8667 |
|  |  | 0.6 | 0.4 | 0.2 | 0.8386 |
|  |  |  |  | 0.3 | 0.849 |
|  |  |  |  | 0.4 | 0.8638 |
|  |  | 0.7 | 0.3 | 0.2 | 0.8618 |
|  |  |  |  | 0.3 | 0.8396 |
|  |  |  |  | 0.4 | 0.87 |

Based on the parameter study, the best results are observed on the unsupervised model using pairwise contrastive loss, which gives an SC of 0.8104. While it is a trivial result that adding labels improves the clustering metric, we chose to include it to observe the difference between the best parameters with labels (supervised) vs. the best parameters when labels are not provided (unsupervised). We observe the drop is 0.0842, which is approximately a drop of 10% when going from the best supervised model (SC = 0.8946) to the best unsupervised model (SC = 0.8104). Using the best parameters from the parameter study (*pt* = 0.6, *nt* = 0.4, *m* = 0.2), the validation dataset embeddings were visualized before and after to observe how the contrastive learning framework contributes to better clustering of the classes.

A 2-dimensional t-SNE plot [48, 49] was used to observe the different embeddings before and after the Siamese Neural Network was run. The reason for using t-SNE is that the local relationships and subtle structures in the data are revealed using t-SNE, which is why it is used on genomic data [50]. In all such figures, x1 and x2 represent the axes of the 2-dimensional plots for ease of observing the visualizations for these multidimensional embeddings. Since the SNN learns pairwise distances and hence new embeddings instead of probabilities, these figures show the shift in embeddings before and after the model learns to bring similar classes together while pushing different classes apart. Figure 5a and Figure 5b show the t-SNE plots of GenSLM embeddings before and after learning. The SC improved from 0.4543 to 0.7309 on the validation set including all variants without being given any labels, which indicates tighter clustering of the embeddings following the SNN training.

**Figure 5:**
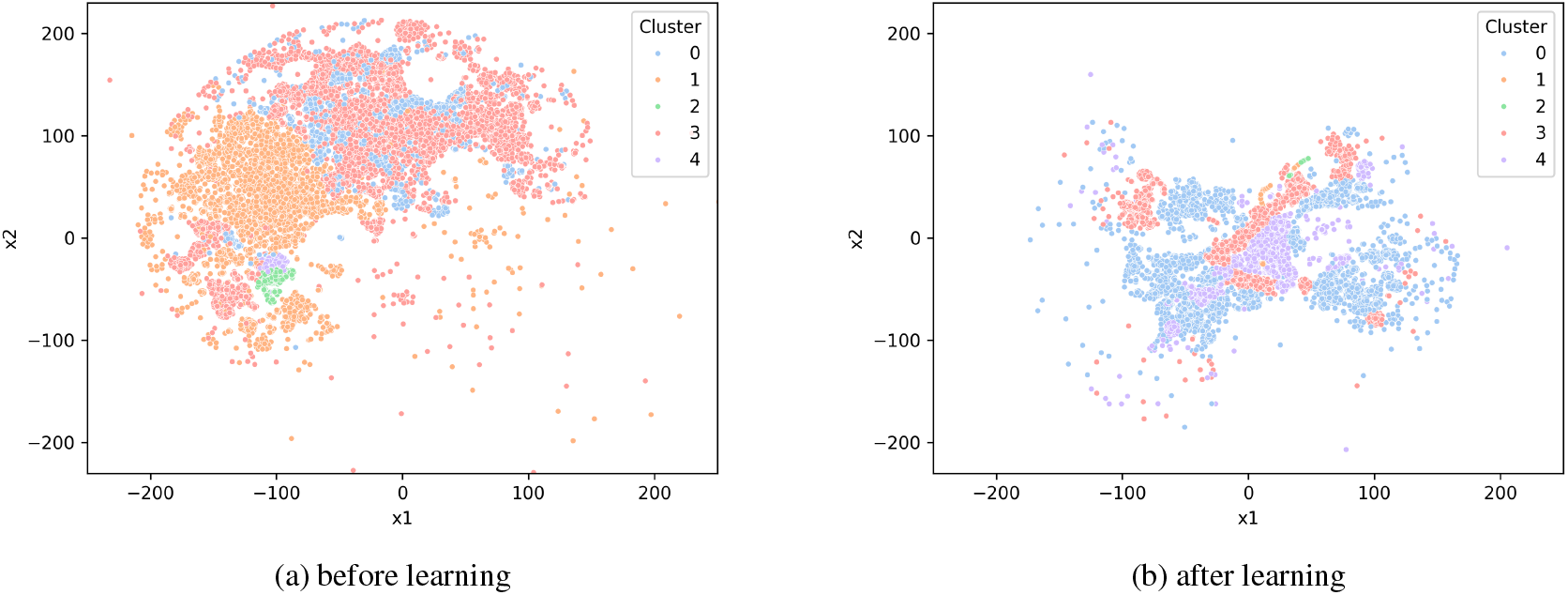
t-SNE embeddings of SARS-CoV-2 variants before and after learning

#### 5.2.2 Zero-shot performance on variants not in train set

While the metrics in the previous section were on the validation set, they included variants that the model previously learned during train time. As a result, the metrics contain a mix of ‘seen’ and ‘unseen’ variants. In this section, the goal is to evaluate the zero-shot capabilities of the SNN. The SNN model by itself only works on sampled pairs of triplets. In SNNs it is common to allow inference by taking only the embedding layer of the trained model and passing the test sequences through it to get the embeddings. However, for quick inference, it is beneficial to have the model work on a set of test embeddings. In order to do this, we chose to add a prediction head to learned SNN embeddings (80%) and allowed it to make predictions on the test (20%) embeddings. Once again, this chronological ordering simulates the scenario of the pandemic where the variants occur over time.

##### Generating embeddings from the SNN

For this step, we use the unsupervised pairwise contrastive model obtained previously based on the parameter study. Due to the scale of the dataset containing all variants, we chose to load the data in batches.

##### Framing the novelty detection problem

Next, we added an extra step where we have the model predict if it is a variant it has ‘seen’ before or whether the variant is ‘unseen’. This creates a new set of labels where we frame the problem as a *novelty detection problem* in computer science to recognize the capabilities of the SNN in identifying variants it has not encountered previously. This is notably a misnomer in the context of this problem since we are not discovering any *virus*, but rather using the framework to identify classes it has not ‘seen’ during train time.

##### Prototype learning

In this step, we apply prototype learning to the dataset in order to obtain the overall distribution of the embeddings. The MiniBatch K-Means clustering algorithm is chosen because of its ability to scale efficiently given the size of the dataset with all variants. For this experiment, we chose *k* = 200 and *batches* = 2048. The cluster centroids from the SNN trained embeddings serve as the prototype vectors for the next step.

##### Novelty scoring based on distance

This step follows distance-based sample scoring similar to the K-Nearest Neighbors algorithm. The Euclidean distance between each embedding and all prototype centroids is calculated, and the minimum distance is the ‘novelty score’ assigned to that embedding. This means that samples belonging to variants previously encountered will be closer to a previously learned ‘prototype’ while samples from variant classes not previously encountered will be farther away. Once again, this step is performed in batches owing to the scale of the dataset.

##### Threshold selection and prediction

Once the novelty scores are obtained on both the train and test embeddings, the 95th percentile of the training scores is considered to obtain the *threshold*. The samples in the test dataset that are greater than this threshold are classified as novel, while those below the threshold are classified as previously seen.

#### 5.2.3 Low-Rank Adaptation (LoRA) adapter

The question we aim to answer here is: *How can domain-specific metric learning improve the prediction of new classes during inference time?*. As a parameter-efficient baseline for comparison, we use low-rank adaptation (LoRA), first proposed by Hu et al. [51]. LoRA can only be used as an adapter in this case and cannot be used as a standalone method since it inherently requires supervision. However, the proposed contrastive learning framework is capable of supporting the task of detecting unseen variants. In this scenario, we use LoRA as an adapter that freezes the original weights and learns a ‘low-rank trainable update’ matrix that adapts the model while fine-tuning. This is done using the following rules:

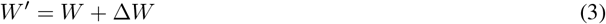

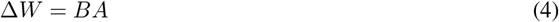

where 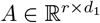 and 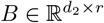, with rank *r << min*(*d*_1_, *d*_2_).

We do this by initializing a custom TensorFlow layer that takes into account the number of output units, rank *r*, and the scaling factor *α* as the parameters. This process is repeated by taking apart the proposed SNN model, replacing all the Dense layers with the LoRA adapters, transferring the pre-trained weights, and stitching it back together as part of the SNN framework. This means *W* is the base matrix with the pre-trained weights while *A* and *B* are the low-rank trainable matrices. Consistent with the model in the previous section, we train the SNN with the pairwise contrastive loss function. We calculate the prototypes with MiniBatch K-means clustering, use a distance-based novelty scoring, and finally select a threshold of the 95th percentile for final novelty prediction as discussed previously. Figure 6 and Figure 7 show the performance of the SNN and SNN + LoRA methods relative to the ground truth. We observe that adding the LoRA adapter did not significantly improve performance, and that the use of the SNN framework on the ESM-2 embeddings is sufficient for the zero-shot variant detection problem.

**Figure 6:**
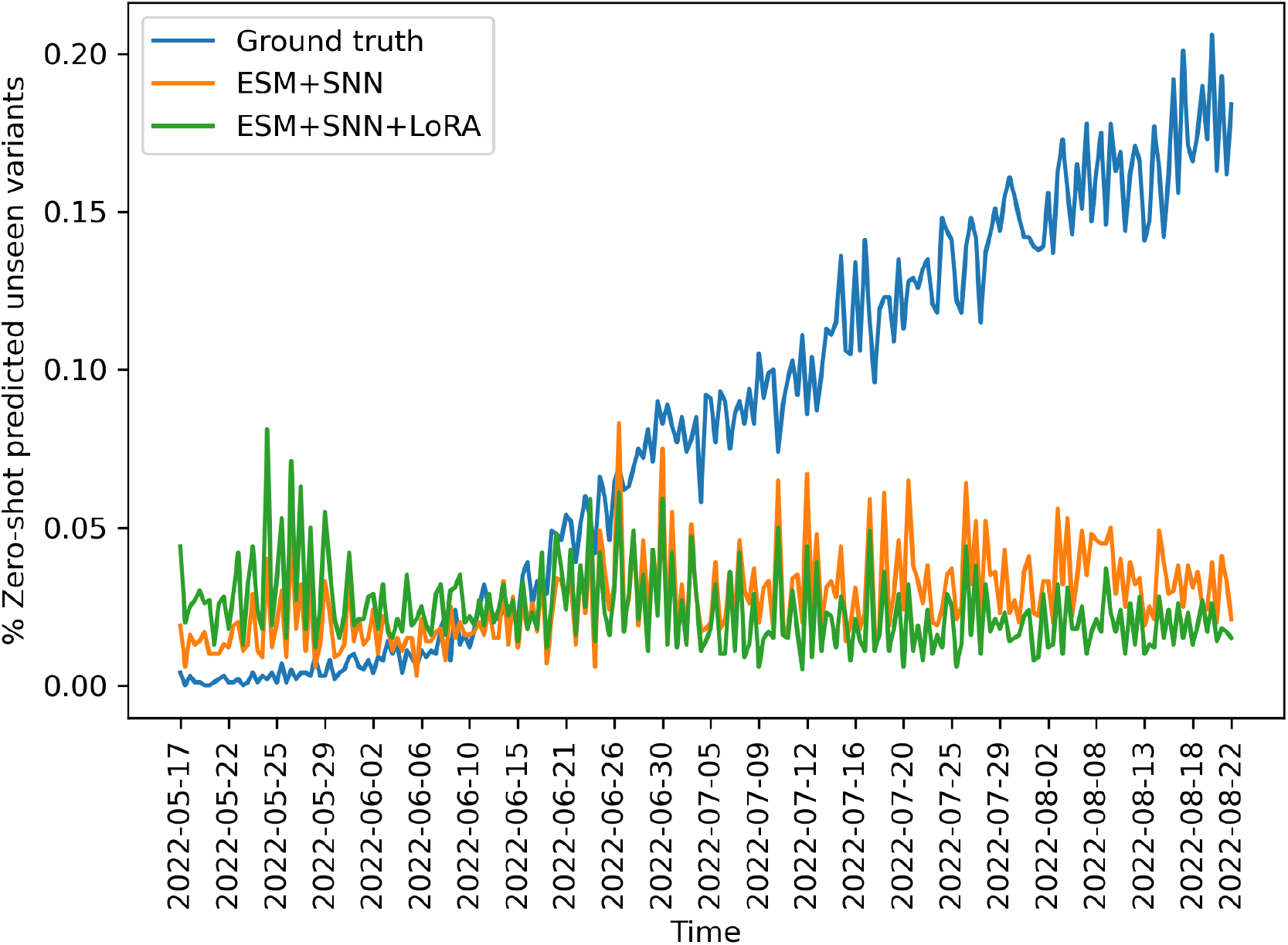
Zero-shot performance of variants over time

**Figure 7:**
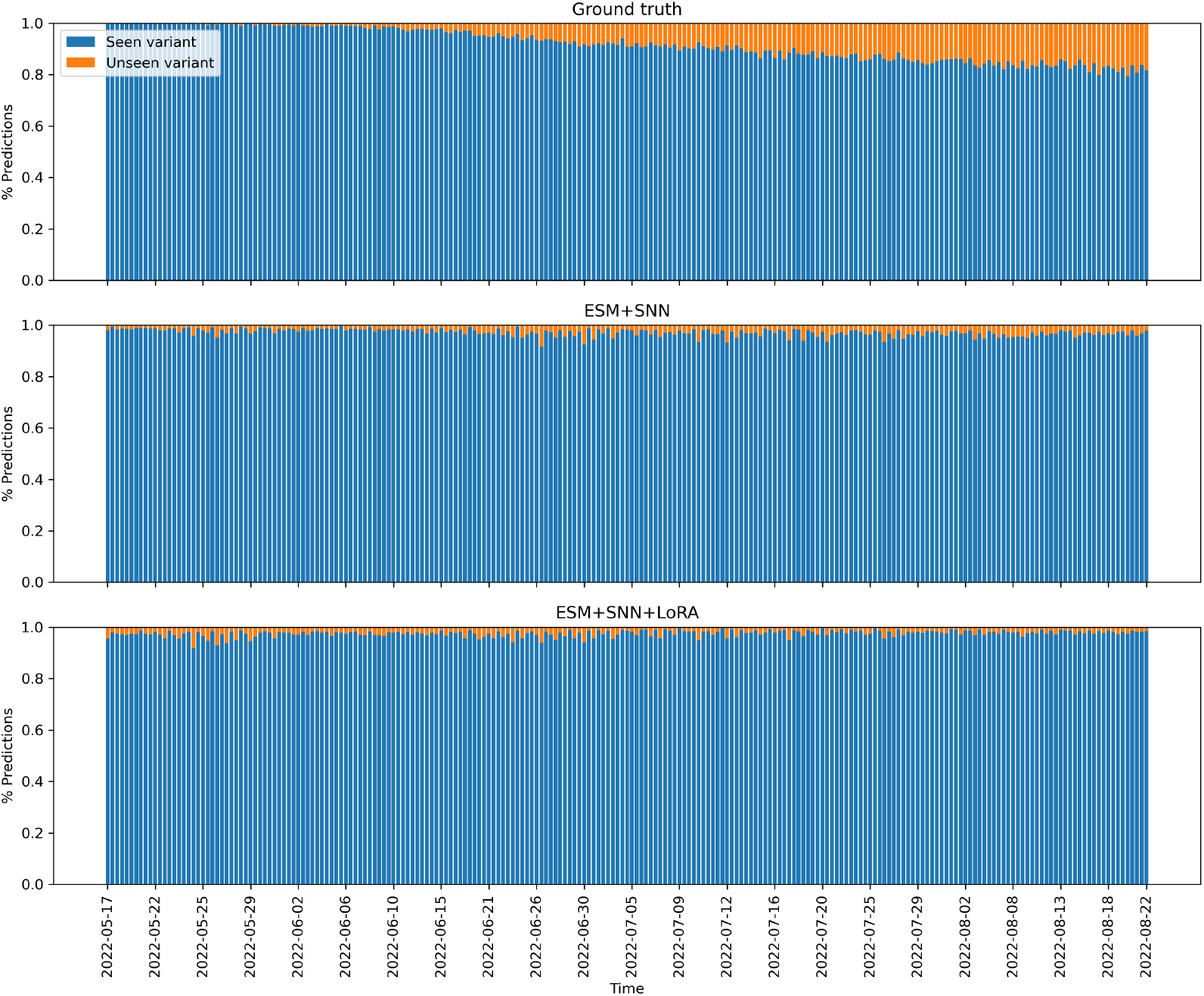
Predictions across different models

#### 5.2.4 Application to detecting emerging variants on the held-out test set

Results from clustering the dataset can be applied to the problem of detecting emerging variants of SARS-CoV-2. In order to prove that there is some benefit to using large language models in this area, prior results from a previous model are replicated, and the proposed LLM-based approach is compared to it. The previously proposed framework by de Hoffer et al. [39] is an unsupervised framework that relies on the Levenshtein distances only as a similarity metric. In this approach, the authors start by computing the Levenshtein distances among all pairs of sequences. Following this, they generate a hierarchical clustering for each timestamp. This generates a dendrogram for each timestamp, and Ward’s method is used to prune the tree and fix the number of clusters. The metric used by the authors is *rW* = 100 which is determined empirically, and this same metric is used in the proposed method for fair comparison. Finally, to ensure that the de Hoffer et al. [39] can be compared to the proposed approach, adjusted rand index (ARI) is used as a metric to compare the cluster alignment of their cluster predictions to the gold standard PANGO labels.

For the proposed framework, the large language model embeddings and the contrastive learning framework are used from the previous section. In the first step, cosine distances are computed among all pairs of LLM embeddings predicted by the SNN with the optimal parameters as determined by the previous parameter study. The LLM used here is also GenSLM as it has shown the best performance among all the other LLMs. In the second step, a hierarchical clustering is generated for each timestamp and Ward’s method is used with *rW* = 100 in de Hoffer’s approach and a threshold of 10.0^*−*10^ for the cosine distance in our approach. In the last step, ARI is used to get a quantitative measure of how well the model predicts clusters compared to the gold standard PANGO labels.

Figure 8 and Figure 9 show boxplots of the two approaches over time. Figure 10 and Figure 11 show the line plots of the ARI relative to the ground truth and optimized clusters respectively. Based on the cluster alignment ARI metric, the proposed LLM-based method better at reducing the distances within the same PANGO variants and increasing distances between different PANGO variants compared to the Levenshtein distance only approach by de Hoffer et al. [39]. The figures show that the proposed LLM-based approach is able to maximize the distances between sequence vectors belonging to different clusters while minimizing the distances between similar PANGO variants. The boxplots show that the LLM-based approach is able to achieve this for a significant number of months in the test dataset compared to the previous approach by de Hoffer et al. [39]. Figures 8 and 9 show the results obtained using ARI on LD-only [39] and the proposed LLM-based approach. A significant improvement is observed for all months of the test dataset, with an average improvement of 0.2 in cluster alignment using the LLM-based approach.

**Figure 8:**
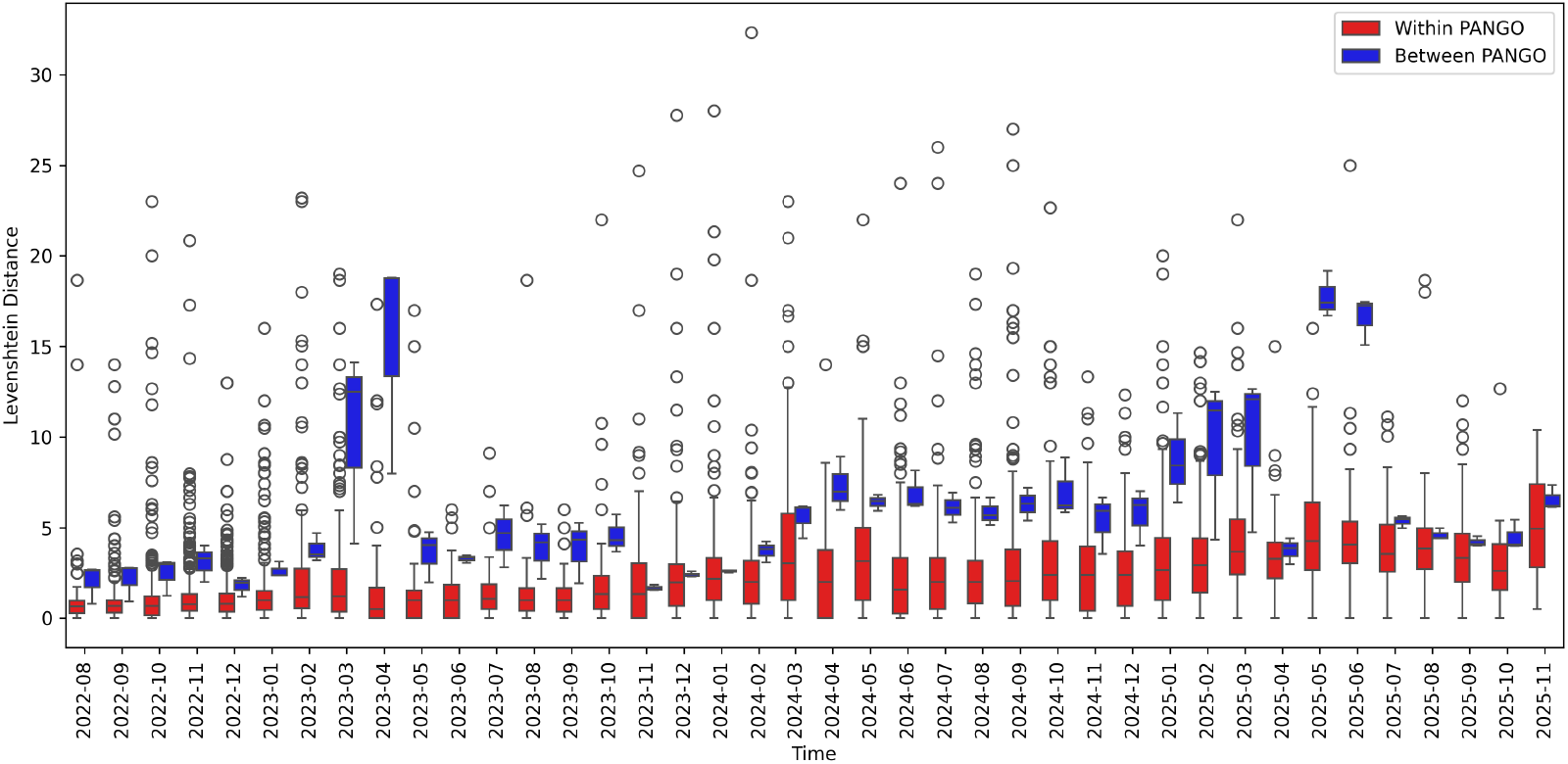
Levenshtein distance metrics over time based on de Hoffer et al. [39]

**Figure 9:**
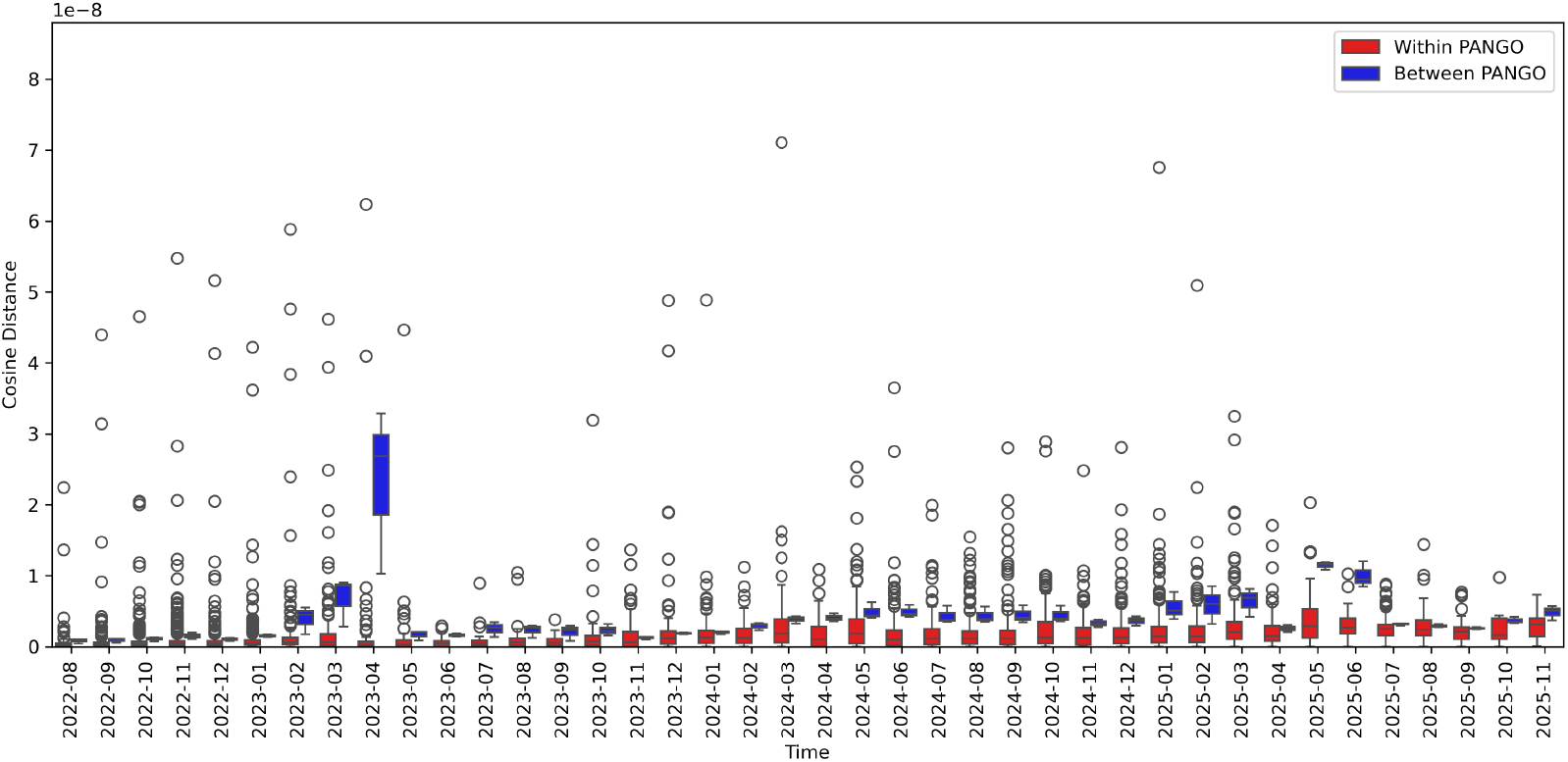
Proposed LLM-based approach metrics using contrastive learning

**Figure 10:**
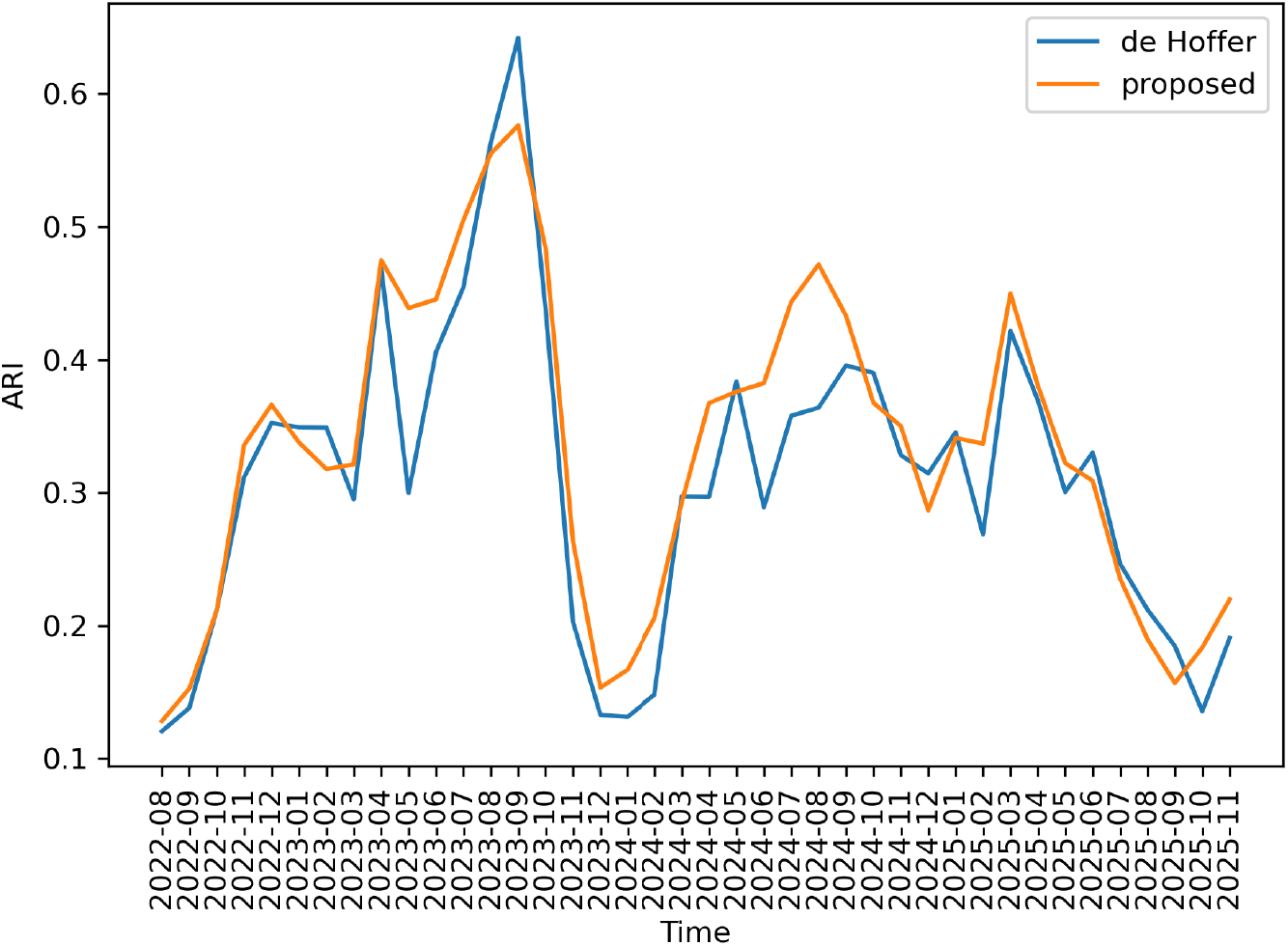
ARI on ground truth clusters

**Figure 11:**
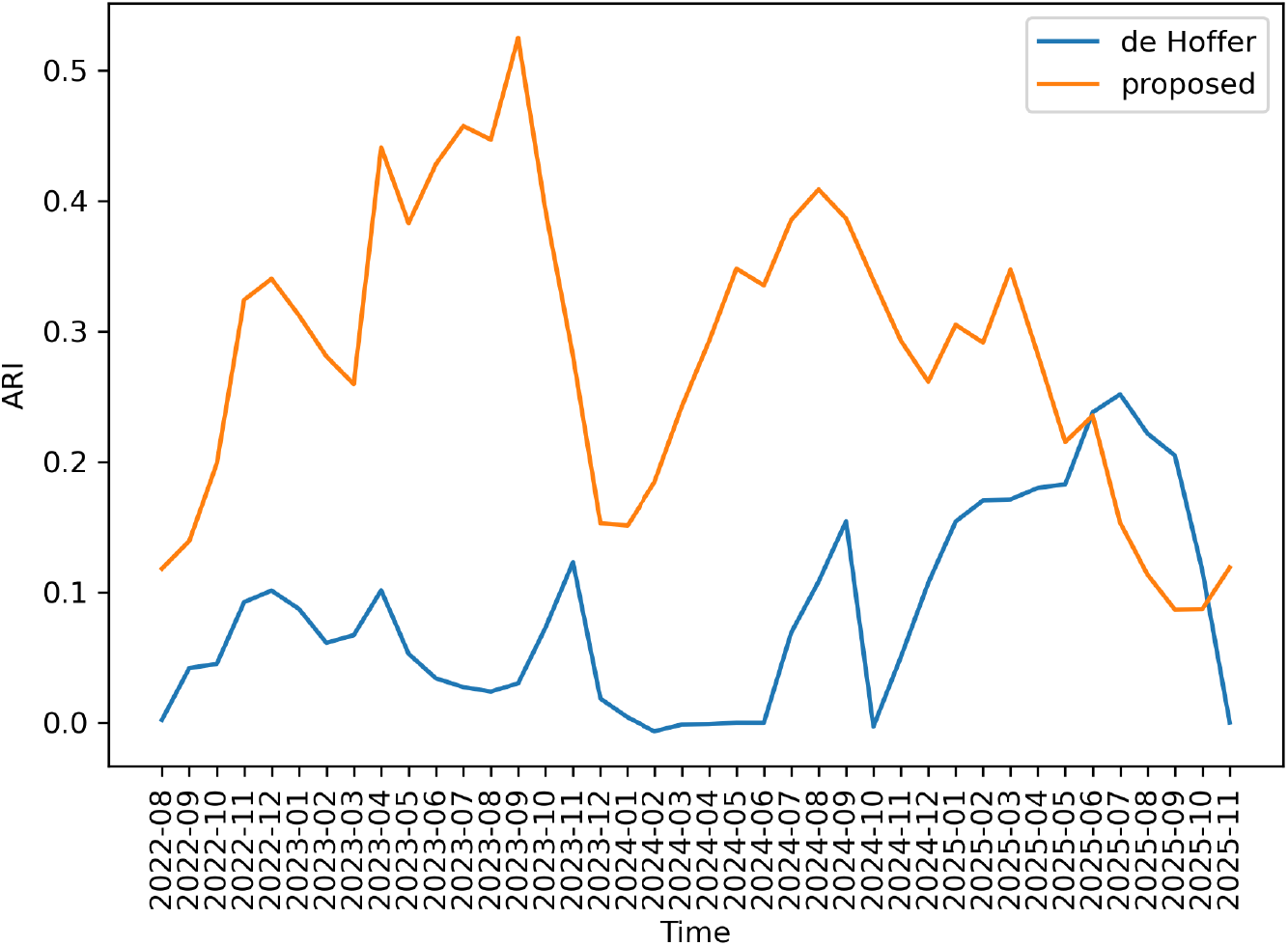
ARI on optimized clusters

## 6 Conclusion

In this paper, we conducted a comparative study of previously proposed language models on SARS-CoV-2 surface glycoprotein (spike) sequences. We discuss how embeddings extracted from previously proposed models perform on classifying and clustering different variants. We then propose using a contrastive learning framework using a Siamese neural network in an unsupervised manner to compare different SARS-CoV-2 protein sequences. The benefit of using an unsupervised contrastive learning framework is that we do not rely on prior labeling methods or embedding vectors alone to compare different protein sequences. Using our framework, we show we can identify that the sequences come from different groups (variants) and that the proposed LLM-based approach shows an improvement of 0.2 in terms of the adjusted rand index clustering on test data compared to a previously proposed approach.

## 7 Code

Code for this paper is publicly available here: https://github.com/sblittlefield/covid-contrast.

## Acknowledgments

We would like to acknowledge Dr. Sergei Maslov from the Bioengineering department at the University of Illinois at Urbana-Champaign for his valuable feedback on our work. This work utilizes the HAL (National Science Foundation’s Major Research Instrumentation program, grant #1725729) and DeltaAI (National Science Foundation, award OAC 2320345 and the State of Illinois) resources from the Advanced Cyberinfrastructure Coordination Ecosystem: Services & Support (ACCESS) program, which is supported by U.S. National Science Foundation grants #2138259, #2138286, #2138307, #2137603, and #2138296. We would also like to acknowledge all laboratories and authors that contributed sequence data used in this paper.

## References

[1] B. Yu and T. Xiaonan, “Comparison of COVID-19 and influenza characteristics,” Journal of Zhejiang University. Science. B, vol. 22, no. 2, p. 87, 2021.

[2] M. A. Johansson, T. M. Quandelacy, S. Kada, P. V. Prasad, M. Steele, J. T. Brooks, R. B. Slayton, M. Biggerstaff, and J. C. Butler, “SARS-CoV-2 transmission from people without COVID-19 symptoms,” JAMA network open, vol. 4, no. 1, pp. e2 035 057–e2 035 057, 2021.

[3] K. Beguir, M. J. Skwark, Y. Fu, T. Pierrot, N. L. Carranza, A. Laterre, I. Kadri, A. Korched, A. U. Lowegard, B. G. Lui et al., “Early computational detection of potential high-risk SARS-CoV-2 variants,” Computers in biology and medicine, vol. 155, p. 106618, 2023.

[4] W. Han, N. Chen, X. Xu, A. Sahil, J. Zhou, Z. Li, H. Zhong, E. Gao, R. Zhang, Y. Wang et al., “Predicting the antigenic evolution of sars-cov-2 with deep learning,” Nature Communications, vol. 14, no. 1, p. 3478, 2023.

[5] B. L. Hie, K. K. Yang, and P. S. Kim, “Evolutionary velocity with protein language models predicts evolutionary dynamics of diverse proteins,” Cell systems, vol. 13, no. 4, pp. 274–285, 2022.

[6] B. Zhou, H. Zhou, X. Zhang, X. Xu, Y. Chai, Z. Zheng, A. C. Kot, and Z. Zhou, “Tempo: A transformer-based mutation prediction framework for sars-cov-2 evolution,” Computers in Biology and Medicine, vol. 152, p. 106264, 2023.

[7] C. B. Jackson, M. Farzan, B. Chen, and H. Choe, “Mechanisms of SARS-CoV-2 entry into cells,” Nature reviews Molecular cell biology, vol. 23, no. 1, pp. 3–20, 2022.

[8] F. Krammer, “The human antibody response to influenza a virus infection and vaccination,” Nature Reviews Immunology, vol. 19, no. 6, pp. 383–397, 2019.

[9] G. Benegas, C. Ye, C. Albors, J. C. Li, and Y. S. Song, “Genomic language models: opportunities and challenges,” Trends in Genetics, 2025.

[10] Z. N. Flamholz, S. J. Biller, and L. Kelly, “Large language models improve annotation of viral proteins,” Research square, 2023.

[11] L.-Y. Wu, Y. Wijesekara, G. J. Piedade, N. Pappas, C. P. Brussaard, and B. E. Dutilh, “Benchmarking bioinformatic virus identification tools using real-world metagenomic data across biomes,” Genome Biology, vol. 25, no. 1, p. 97, 2024.

[12] I. Aksamentov, C. Roemer, E. B. Hodcroft, and R. A. Neher, “Nextclade: clade assignment, mutation calling and quality control for viral genomes,” Journal of open source software, vol. 6, no. 67, p. 3773, 2021.

[13] J. Huddleston, J. Hadfield, T. R. Sibley, J. Lee, K. Fay, M. Ilcisin, E. Harkins, T. Bedford, R. A. Neher, and E. B. Hodcroft, “Augur: a bioinformatics toolkit for phylogenetic analyses of human pathogens,” Journal of open source software, vol. 6, no. 57, 2021.

[14] J. Hadfield, C. Megill, S. M. Bell, J. Huddleston, B. Potter, C. Callender, P. Sagulenko, T. Bedford, and R. A. Neher, “Nextstrain: real-time tracking of pathogen evolution,” Bioinformatics, vol. 34, no. 23, pp. 4121–4123, 2018.

[15] S. Ali, B. Sahoo, A. Zelikovsky, P.-Y. Chen, and M. Patterson, “Benchmarking machine learning robustness in COVID-19 genome sequence classification,” Scientific Reports, vol. 13, no. 1, p. 4154, 2023.

[16] S. Basu, “A study of the dynamics and genetics of COVID-19 through machine learning,” Master’s thesis, University of Illinois Urbana-Champaign, 2020.

[17] G. S. Randhawa, M. P. Soltysiak, H. El Roz, C. P. de Souza, K. A. Hill, and L. Kari, “Machine learning using intrinsic genomic signatures for rapid classification of novel pathogens: COVID-19 case study,” Plos one, vol. 15, no. 4, p. e0232391, 2020.

[18] L. Brierley and A. Fowler, “Predicting the animal hosts of coronaviruses from compositional biases of spike protein and whole genome sequences through machine learning,” PLoS Pathogens, vol. 17, no. 4, p. e1009149, 2021.

[19] S. Basu and R. H. Campbell, “Classifying COVID-19 variants based on genetic sequences using deep learning models,” in System Dependability and Analytics: Approaching System Dependability from Data, System and Analytics Perspectives. Springer, 2022, pp. 347–360.

[20] E. Asgari and M. R. Mofrad, “Continuous distributed representation of biological sequences for deep proteomics and genomics,” PloS one, vol. 10, no. 11, p. e0141287, 2015.

[21] N. Brandes, D. Ofer, Y. Peleg, N. Rappoport, and M. Linial, “ProteinBERT: a universal deep-learning model of protein sequence and function,” Bioinformatics, vol. 38, no. 8, pp. 2102–2110, 2022.

[22] T. Mikolov, I. Sutskever, K. Chen, G. S. Corrado, and J. Dean, “Distributed representations of words and phrases and their compositionality,” Advances in neural information processing systems, vol. 26, 2013.

[23] J. Devlin, M.-W. Chang, K. Lee, and K. Toutanova, “BERT: Pre-training of Deep Bidirectional Transformers for Language Understanding,” in Proceedings of the 2019 Conference of the North American Chapter of the Association for Computational Linguistics: Human Language Technologies, Volume 1 (Long and Short Papers), 2019, pp. 4171–4186.

[24] A. Vaswani, N. Shazeer, N. Parmar, J. Uszkoreit, L. Jones, A. N. Gomez, Ł. Kaiser, and I. Polosukhin, “Attention is all you need,” Advances in neural information processing systems, vol. 30, 2017.

[25] A. Radford, K. Narasimhan, T. Salimans, I. Sutskever et al., “Improving language understanding by generative pre-training,” OpenAI, 2018.

[26] J. Ito, A. Strange, W. Liu, G. Joas, S. Lytras, and K. Sato, “A protein language model for exploring viral fitness landscapes,” Nature communications, vol. 16, no. 1, p. 4236, 2025.

[27] N. N. Thadani, S. Gurev, P. Notin, N. Youssef, N. J. Rollins, D. Ritter, C. Sander, Y. Gal, and D. S. Marks, “Learning from prepandemic data to forecast viral escape,” Nature, vol. 622, no. 7984, pp. 818–825, 2023.

[28] Z. Lin, H. Akin, R. Rao, B. Hie, Z. Zhu, W. Lu, N. Smetanin, A. dos Santos Costa, M. Fazel-Zarandi, T. Sercu, S. Candido et al., “Language models of protein sequences at the scale of evolution enable accurate structure prediction,” bioRxiv, 2022.

[29] R. M. Rao, J. Meier, T. Sercu, S. Ovchinnikov, and A. Rives, “Transformer protein language models are unsupervised structure learners,” bioRxiv, 2020. [Online]. Available: https://www.biorxiv.org/content/10.1101/2020.12.15.422761v1

[30] M. Zvyagin, A. Brace, K. Hippe, Y. Deng, B. Zhang, C. O. Bohorquez, A. Clyde, B. Kale, D. Perez-Rivera, H. Ma et al., “GenSLMs: Genome-scale language models reveal SARS-CoV-2 evolutionary dynamics,” The International Journal of High Performance Computing Applications, vol. 37, no. 6, pp. 683–705, 2023.

[31] E. Nguyen, M. Poli, M. Faizi, A. Thomas, M. Wornow, C. Birch-Sykes, S. Massaroli, A. Patel, C. Rabideau, Y. Bengio et al., “Hyenadna: Long-range genomic sequence modeling at single nucleotide resolution,” Advances in neural information processing systems, vol. 36, pp. 43 177–43 201, 2023.

[32] A. Elnaggar, M. Heinzinger, C. Dallago, G. Rehawi, Y. Wang, L. Jones, T. Gibbs, T. Feher, C. Angerer, M. Steinegger et al., “Prottrans: toward understanding the language of life through self-supervised learning,” IEEE transactions on pattern analysis and machine intelligence, vol. 44, no. 10, pp. 7112–7127, 2021.

[33] A. Elnaggar, H. Essam, W. Salah-Eldin, W. Moustafa, M. Elkerdawy, C. Rochereau, and B. Rost, “Ankh: Optimized protein language model unlocks general-purpose modelling,” arXiv preprint arXiv:2301.06568, 2023.

[34] G. Brixi, M. G. Durrant, J. Ku, M. Naghipourfar, M. Poli, G. Sun, G. Brockman, D. Chang, A. Fanton, G. A. Gonzalez et al., “Genome modelling and design across all domains of life with evo 2,” Nature, pp. 1–13, 2026.

[35] G. Koch, R. Zemel, R. Salakhutdinov et al., “Siamese neural networks for one-shot image recognition,” in ICML deep learning workshop, vol. 2, no. 1. Lille, 2015.

[36] S. Madan, V. Demina, M. Stapf, O. Ernst, and H. Fröhlich, “Accurate prediction of virus-host protein-protein interactions via a Siamese neural network using deep protein sequence embeddings,” Patterns, vol. 3, no. 9, 2022.

[37] S. Tsukiyama, M. M. Hasan, S. Fujii, and H. Kurata, “LSTM-PHV: prediction of human-virus protein–protein interactions by LSTM with word2vec,” Briefings in bioinformatics, vol. 22, no. 6, p. bbab228, 2021.

[38] S. Basu, R. H. Campbell, and K. Karahalios, “Detection of Novel COVID-19 Variants with Zero-Shot Learning,” HealthNLP’23, IEEE ICHI, 2023.

[39] A. de Hoffer, S. Vatani, C. Cot, G. Cacciapaglia, M. L. Chiusano, A. Cimarelli, F. Conventi, A. Giannini, S. Hohenegger, and F. Sannino, “Variant-driven early warning via unsupervised machine learning analysis of spike protein mutations for covid-19,” Scientific Reports, vol. 12, no. 1, p. 9275, 2022.

[40] A. Rambaut, E. C. Holmes, Á. O’Toole, V. Hill, J. T. McCrone, C. Ruis, L. du Plessis, and O. G. Pybus, “A dynamic nomenclature proposal for SARS-CoV-2 lineages to assist genomic epidemiology,” Nature microbiology, vol. 5, no. 11, pp. 1403–1407, 2020.

[41] Á. O’Toole, E. Scher, A. Underwood, B. Jackson, V. Hill, J. T. McCrone, R. Colquhoun, C. Ruis, K. Abu-Dahab, B. Taylor et al., “Assignment of epidemiological lineages in an emerging pandemic using the pangolin tool,” Virus evolution, vol. 7, no. 2, p. veab064, 2021.

[42] J. Bromley, I. Guyon, Y. LeCun, E. Säckinger, and R. Shah, “Signature verification using a “siamese” time delay neural network,” Advances in neural information processing systems, vol. 6, 1993.

[43] K. Kieft and K. Anantharaman, “Virus genomics: what is being overlooked?” Current opinion in virology, vol. 53, p. 101200, 2022.

[44] D. P. Kingma and J. Ba, “Adam: A method for stochastic optimization,” arXiv preprint arXiv:1412.6980, 2014.

[45] V. Kindratenko, D. Mu, Y. Zhan, J. Maloney, S. H. Hashemi, B. Rabe, K. Xu, R. Campbell, J. Peng, and W. Gropp, “HAL: Computer system for scalable deep learning,” in Practice and experience in advanced research computing, 2020, pp. 41–48.

[46] B. Bode, G. Bauer, L. Herriott, V. Kindratenko, and W. Gropp, “DeltaAI: A National Resource for AI/ML Research,” in Practice and Experience in Advanced Research Computing 2025: The Power of Collaboration, 2025, pp. 1–4.

[47] T. J. Boerner, S. Deems, T. R. Furlani, S. L. Knuth, and J. Towns, “Access: Advancing innovation: NSF’s advanced cyberinfrastructure coordination ecosystem: Services & support,” in Practice and experience in advanced research computing 2023: Computing for the common good, 2023, pp. 173–176.

[48] G. E. Hinton and S. Roweis, “Stochastic neighbor embedding,” Advances in neural information processing systems, vol. 15, 2002.

[49] L. Van der Maaten and G. Hinton, “Visualizing data using t-SNE,” Journal of machine learning research, vol. 9, no. 11, 2008.

[50] W. Li, J. E. Cerise, Y. Yang, and H. Han, “Application of t-SNE to human genetic data,” Journal of bioinformatics and computational biology, vol. 15, no. 04, p. 1750017, 2017.

[51] E. J. Hu, Y. Shen, P. Wallis, Z. Allen-Zhu, Y. Li, S. Wang, L. Wang, W. Chen et al., “Lora: Low-rank adaptation of large language models.” ICLR, vol. 1, no. 2, p. 3, 2022.

